# Homeostatic changes maintain the gain control of spinal motoneurones across the lifetime of C57BL/6J mice

**DOI:** 10.1101/2022.05.19.492717

**Authors:** S Goltash, MH Jensen, KP Dimintiyanova, DB Jensen, J Lehnhoff, M Moldovan, CF Meehan

**Author notes:** Correspondence to: Claire F. Meehan, Department of Neuroscience, University of Copenhagen, Panum Institute, 2200 Copenhagen N, Denmark. **Author contributions (using CRediT roles)** SG: Conceptualization; Investigation; Formal analysis; Writing - original draft MHJ: Conceptualization; Investigation: Formal analysis; Writing - review & editing DBJ: Formal analysis; Writing - review & editing KPD: Formal analysis; Visualization; Writing - original draft JL: Investigation; Supervision; Writing - review & editing MM: Resources. Writing - review & editing CFM: Funding acquisition; Conceptualization; Investigation, Data curation; Visualization; Supervision; Writing - original draft.

## Abstract

Age-related changes in the excitability of spinal motoneurone have been observed in mouse models of neurodegenerative diseases affecting these neurones. How the excitability of spinal motoneurones change with healthy ageing in mice and how this compares with that seen in neurodegenerative diseases is unknown. Therefore, we performed *in vivo* intracellular recording from identified spinal motoneurones in C57BL/6 mice at three different ages (100, 300-400 and 600-750 days old). Behavioral tests confirmed a linear reduction in motor function across these ages (using the rotorod test).

Significant differences were observed with respect to the features of individual somatic action potential with ageing including a decreased rate of rise and fall in aged mice. Surprisingly, the rate of rise of the action potential at the initial segment was altered in middle aged mice. Immunohistochemical labelling of the axon initial segment of the motoneurones confirmed structural changes occurring at middle age (decreased length and diameter) but returning to the earlier parameters in aged mice. To explore the effects on repetitive firing, this was tested across the age groups which showed surprising little difference as the mice aged, with a similar rheobase and I-f gain across all age groups (with the exception of a lower voltage threshold for action potential initiation in middle-aged mice). However, amplitudes of the after-hyperpolarization and the input resistance were both found to be significantly altered with age.

We conclude that there are changes occurring in the intrinsic properties of spinal motoneurones that control their excitability over the lifetime of mice, although these do not develop in a linear fashion from young to old. We propose that these changes are homeostatic in nature and are able to compensate for one another to maintain a constant gain control across the lifetime.

## INTRODUCTION

Abnormal neuronal excitability has been demonstrated in several neurodegenerative diseases affecting the motor system. Despite ageing being one of the biggest risk factors for the development of such diseases, we still do not fully understand the excitability changes that occur in spinal motoneurones during the normal ageing process. Consequently, it is crucial to determine the extent to which changes observed in disease models represent either an accelerated ageing process or distinctly different pathological processes. Furthermore, even normal ageing leads to profound impairments in motor function. Explanations for this have tended to focus on sarcopenia or the loss of motoneurones and so the degree to which functional changes in motoneurone excitability may also contribute to the normal age-related decline in motor function is not clear.

Given the relative inaccessibility of the spinal motoneurones for direct electrophysiological investigation in humans, it is difficult to assess how motoneurone excitability changes across our lifetime. Reductions in the Hoffman reflex (an electrophysiological analog of the stretch reflex) would suggest either a reduction in the intrinsic excitability of the spinal motoneurone or a change in synaptic release at the Ia afferent - motoneurone synapse (Schmidt *et al*., 1982;Scaglioni *et al*., 2002;Sabbahi & Sedgwick, 1982). Paradoxically however, post-activation depression of the Ia EPSP appears to be reduced in ageing (Hedegaard *et al*., 2015). A reduction in the firing rate of motor units is also observed (Hassan *et al*., 2021 Orssatto *et al*., 2021, 2022), although again, the extent to which this reflects a reduction in the synaptic drive to the spinal motoneurones or a decrease in their intrinsic responsiveness is difficult to determine from electromyographical (EMG) recordings.

Intracellular recording experiments in animals provide a method to directly test motoneurone excitability at different ages. Surprisingly, only a small number of studies have recorded from motoneurones in aged animals, these being cats and rats. In both species, ageing was associated with increases in input resistance (Kalmar *et al*., 2009;Morales *et al*., 1987), consistent with observations of reductions in motoneurone soma size (Liu *et al*., 1996). Decreases in the rheobase currents for single action potentials have, also, been observed in both species (Kalmar *et al*., 2009;Morales *et al*., 1987). These features are all surprisingly more consistent with an increase in intrinsic excitability. This suggests that rather than contributing to motor decline, homeostatic processes may act to increase motoneurone excitability to combat other changes occurring. These electrophysiological parameters are concerned with single action potentials but for maximal force generation, repetitive firing of motoneurones at high frequencies is necessary for tetanic contractions in muscle fibers (Jensen *et al*., 2018). Surprisingly, features of repetitive firing have only been investigated once in aged animals (in rat) which showed a decreased gain (I-f gain) despite increases in calcium persistent inward currents (Kalmar *et al*., 2009).

Increased motoneurone excitability has been previously described in mouse models of age-related neurodegenerative diseases affecting the motor system, such as Amyotrophic Lateral Sclerosis (Bonnevie *et al*., 2020; Delestree *et al*., 2014; Huh *et al*., 2020; Jensen *et al*., 2020a; Jørgensen *et al*., 2021; Meehan *et al*., 2010a). It has been suggested that the degenerative process in these motoneurones may arise from a homeostatic dysregulation of excitability (Kuo *et al*., 2020). But no studies using intracellular recording have measured excitability changes with normal ageing in mice to investigate normal homeostasis with healthy ageing which is imperative to allow comparison with the changes that have been observed in mice models of neurodegenerative disorders.

Neuronal excitability itself is heavily controlled by specialized highly excitable domains including, most importantly, the axon initial segment (AIS) - a region of high density clustering of Na^+^ channels forming the site of action potential initiation in most neurones. Plasticity in the size and location of the AIS has been established as a highly effective homeostatic mechanism by which neurones can alter their intrinsic excitability in response to changes in input (Kuba *et al*., 2010) and activity levels (Grubb & Burrone, 2010). Plasticity of motoneurone AISs have already been observed following peripheral nerve injury (Meehan *et al* 2011), intramuscular Botulinum toxin injections (Jensen *et al*., 2020b) and spinal cord injury (Azam *et al*., 2014). Importantly, we have also shown changes the length and diameters of motoneurone AISs in Amyotrophic Lateral Sclerosis (Bonnevie *et al*., 2020; Jørgensen *et al*., 2021). AIS plasticity would therefore be a highly effective way to maintain motoneurone output during age-related perturbation in motor inputs.

The aim of the current study was therefore to investigate the effect of ageing on the intrinsic excitability and axon initial segments of mice spinal motoneurones to provide not only an understanding of the ageing process of spinal motoneurones, but also a background for comparison of changes observed in mouse models of age-related neurodegenerative disease.

## METHODS

All experiments were performed in accordance with EU regulations (Council Directive 2010/63/EU). The protocols used were approved by the Danish National Animal Experiment Committee (2015-15-0201-00545).

### Mice

23 C57BL/6J male mice (Janvier, France) were used for electrophysiology experiments and 27 mice were used for anatomy. They were categorized into 3 different age groups of 100-200 days old (young adults, YA), 300-400 days old (middle-aged; MA) and 600-750 days old (aged; A). Different mice were used for the anatomy and electrophysiology experiments to control for any possible influence of prolonged anaesthesia, immobilization, and nerve stimulation (occurring during electrophysiology experiments) on AIS geometry. Mice used for behavioural (RotaRod) tests were also used for either the electrophysiological or anatomical experiments.

### Rotarod test

To assess general motor function at three different ages investigated, the mice were placed on an Ugo Basile 7650 accelerating RotaRod (Ugo Basile Srl, 21,024 Comerio VA, Italy). The counter was then started and the rod was accelerated from 4 to 40 rpm over a period of 300s. The RotaRod endurance time was calculated as the mean endurance time of 3 consecutive measurements repeated at 10 minutes intervals. Trials were excluded if the mice held on to the rod instead of walking or voluntarily jumped off, in which case the mouse was rested for 10 minutes and the trial then repeated.

### Electrophysiology experiments

Mice were initially anaesthetized with inhaled isoflurane followed by an intraperitoneal (IP) injection of a mixture of ketamine and xylazine (0.3 ml ketamine in 4.7 ml saline and 0.5 ml xylazine in 4.5 ml saline mixed together, induction dose of 0.1 mL/10 g). Maintenance of anaesthesia was achieved with top-up doses of ketamine/xylazine 0.05 ml/10 g every 30 minutes throughout the experiment, delivered via intraperitoneal cannulas. Adequacy of anaesthesia was assessed by lack of reflexes to a brief noxious pinch of the hind limb. All mice were administered a single dose of atropine (0.02mg) at the start of the surgical procedures. A rectal probe measured the core body temperature and maintained it at approximately 37°C by a thermostatically controlled heat pad below and heat lamp above the mice.

Surgical procedures for the electrophysiological experiments were performed as previously described (Meehan et al. 2010). Briefly, IP cannulae were inserted for anaesthetic delivery and a tracheotomy tube was inserted into the trachea to allow for subsequent ventilation. The left sciatic nerve was dissected into the tibial and common peroneal branches and a hemi-laminectomy was performed on the L1 vertebra allowing access to the caudal L3-rostral L4 spinal cord segments.

Mice were placed in a modified Narishige stereotactic frame with the head supported by a head holder and vertebral clamps placed on vertebrate T12 and L2. The skin around the hind leg dissection and the spinal cord were tied to the frame to create paraffin oil pools (with pool temperatures maintained at approximately 37°C). Once mice were ventilated (70 breaths/min and a tidal volume of approximately 0.2 mL), the neuromuscular blocking agent Pavulon (Pancuronium bromide) was administered (IP diluted 1:10 in saline, administered as an initial dose of 0.2 mL followed by a dose of 0.1 mL once per hour). Under neuromuscular blockade the electrocardiogram (ECG) was monitored continuously and anaesthesia maintained with the same dosages as previously described (surgical level of anaesthesia) and in the event of any increases in heart rate an additional top up was given (although this was rarely necessary).

Two pairs of custom made bipolar stimulating electrodes were placed on the dissected nerves. The incoming volley from the stimulated peripheral nerve was recorded by a silver recording electrode placed on the dorsal-lateral surface of the exposed spinal cord (recorded as cord dorsum potentials). This was initially used to confirm the correct spinal segment and later for timing during antidromic identification of motoneurones. The dura mater of the spinal cord at this level was gently removed with forceps and a glass microelectrode (optimal resistance: 20-30 MΩ) filled with 2M potassium acetate was inserted into the spinal cord operated by an electronic micro-drive. An Axoclamp 2B amplifier was used to record and amplify the changes in membrane potential (Vm) by 10X in either bridge - or discontinuous current clamp (DCC) mode. Outputs were further amplified and then filtered (using amplifiers/filters from Digitimer, UK) and converted from analogue to digital signals using the 1401 analog-to-digital converter, then recorded and analysed using Spike2 software (both Cambridge Electronic Design (CED), Cambridge, UK).

#### Measurement of intrinsic properties of the motoneurones

Motoneurones were identified by the timing of all-or-none antidromic spikes relative to arrival of the cord dorsum potential. Analyses of features of single action potentials were made from averages of successive action potentials (minimum of 10 spikes) and the spike height and width at 2/3 amplitude were measured. To measure the distance between the initial segment (IS) and somatodendritic (SD) components of the action potential, the antidromic spike was differentiated (dv/dt) revealing two peaks representing the maximum rate of rise of the action potential as it arrives at the initial segment and then the soma (Fig.2D). The difference in time between two peaks was also measured. The later trough in the differentiated action potential indicated the maximum rate of fall of the somatic action potential. All above mentioned features are affected by the Vm, therefore this was measured at the point at which the average was made and confirmed extracellularly. Only cells with a Vm more hyperpolarized than −50 mV with overshooting action potentials were accepted and, where necessary, analyses were performed with respect to Vm using linear regression.

**Figure 1.**
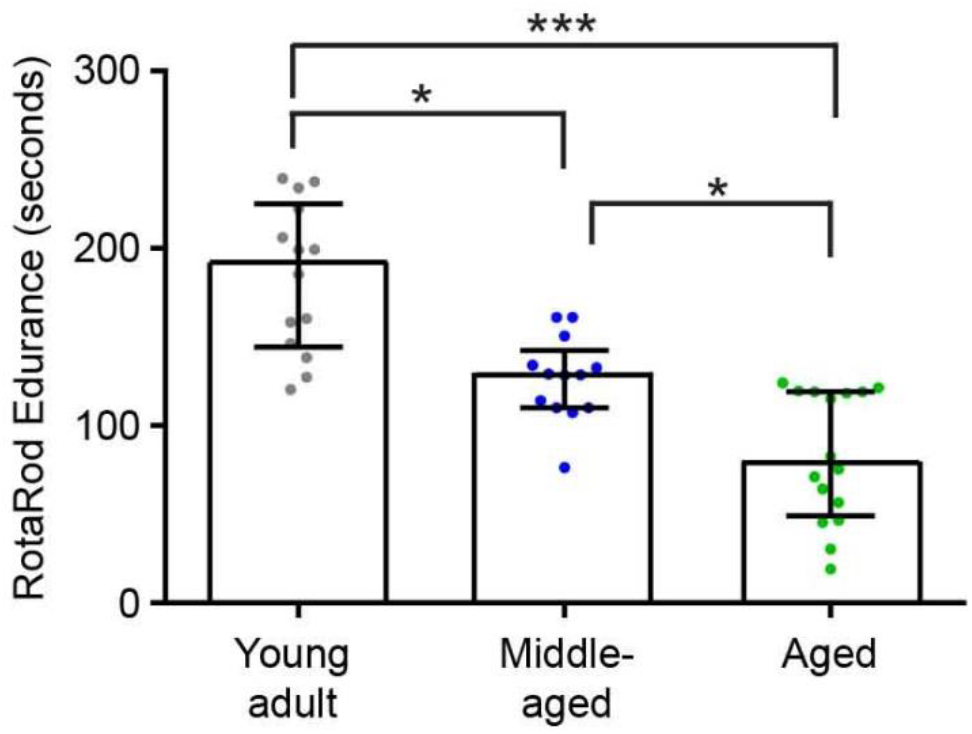
Results from the RotaRod endurance test. Motor performance of young adult (n=14), middleaged (n=13) and aged (n=16) mice was assessed using accelerating RotaRod. Data presented as individual data points with median (+interquartile ranges), *P<0.05, ***P<0.005.

**Figure 2.**
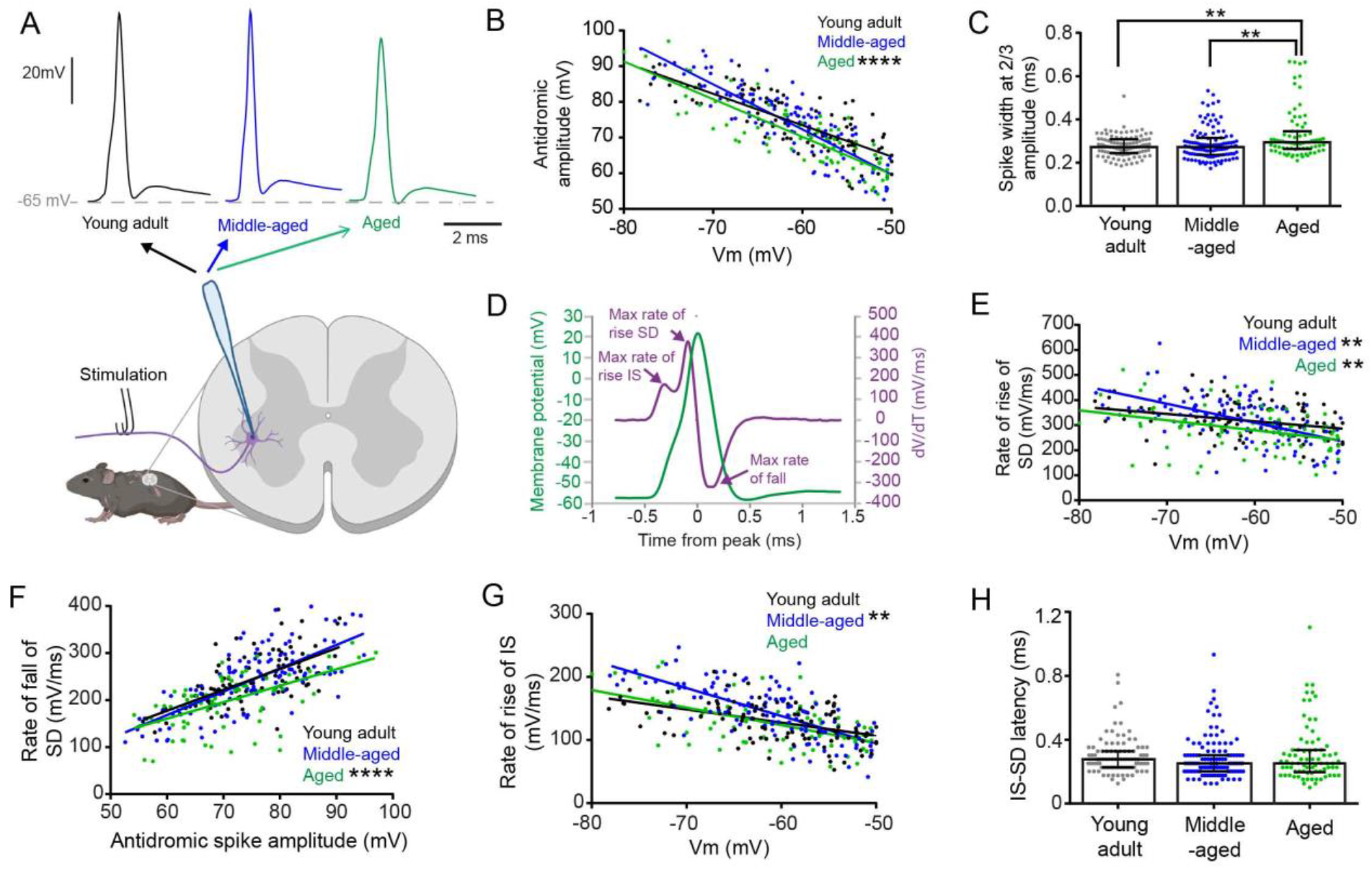
Features of antidromic action potentials in Young Adult (YA, black), Middle-Aged (MA, blue) and Aged (A, green) mice. A. Examples of averaged antidromic action potentials evoked from stimulation of the sciatic nerve recorded intracellularly in motoneurones from YA (black), MA (blue) and Aged (green) mice. B. Graph showing data points for the amplitudes of individual soma-dendritic action potentials by membrane potential. Linear regression lines show the significant relationship between action potential amplitude and resting Vm for motoneurones from YA, MA and aged mice. The angle of the regression slope was significantly steeper in MA mice than both YA and aged mice. The elevation of the regression slope is significantly lower in aged mice than in YA. C. Scattered dot plot of antidromic spike width measured at 2/3 amplitude. The action potentials in the aged mice are significantly wider than action potentials from both YA and MA mice. Data presented as individual data points (cells) with median (+interquartile ranges). D. Example of an average of antidromic action potentials (green) from a motoneurone in an aged mouse with its first derivative (purple) showing two peaks corresponding to the maximal rate of rise of the IS and SD components of the antidromic action potential and a trough corresponding to the maximum rate of fall. E. Graph showing data points for individual action potentials and linear regression lines showing the relationship between the maximum rate of rise of the soma-dendritic (SD) component of the action potential and Vm for motoneurones from YA, MA and aged mice. The angle of the regression slope is significantly steeper in MA mice than both YA and aged mice. The elevation of the regression slope is significantly lower in aged mice than in YA. F. Data points for individual action potentials and linear regression lines showing the relationship between the maximum rate of rise of the IS component of the action potential and Vm for motoneurones from YA, MA and aged mice. The angle of the regression slope was significantly steeper in MA mice than both YA and aged mice. G. Data points for individual action potentials and linear regression lines showing the relationship between the maximum rate of fall of the SD component of the action potential and antidromic spike height in motoneurones from YA, MA and aged mice. The elevation of the regression slope is significantly lower in aged mice than in YA mice. H. Scatter dot plot showing data points for individual cells for IS-SD latency for motoneurones, measured as the time interval between the maximum rate of rise of the IS and the SD components of the antidromic action potential. Lines shown with median (+interquartile ranges) which was not significantly different between groups. For all analysis n= 101 cells YA, 116 cells MA and 77 cells in aged mice. *P<0.05, **P<0.01, ***P<0.001 and ****P<0.0001. Detail of statistical test can be found in tables 2 and 3.

To measure the post-spike after-hyperpolarization (AHP), a short depolarizing current pulse of 1ms was applied through the microelectrode to evoke a single action potential (in bridge mode). The AHP was measured from averages using the decay time to 2/3 of the amplitude of the AHP as the exact return to baseline can often be difficult to assess accurately.

To evoke repetitive firing, triangular depolarizing-repolarizing current ramps were injected through the microelectrode into the cell body and the voltage response was measured using DCC mode (using 3kHz as this appears to be more optimal to evoke repetitive firing and at 8kHz it is often difficult to reliably compensate for capacitance with high resistance electrodes). The instantaneous firing frequencies were calculated in Spike2 and x-y plots exported to Excel (Microsoft Office) to measure the current-frequency gain during the stable primary range or firing.

At the end of the electrophysiology experiments the mice were humanely sacrificed with an overdose of anaesthetic.

### Anatomical experiments

For these experiments mice were euthanized with sodium pentobarbital (130mg/kg), and perfused transcardially with 4% paraformaldehyde diluted in phosphate buffer (pH= 7.4). The spinal cords were removed and the L3-L4 ventral roots were dissected from the 2% fixed tissue for node of Ranvier labeling. The spinal cords from the 4% fixed tissue was post-fixed in 4% paraformaldehyde for an additional 2-3 hours for AIS labelling experiments and then immersed in 30% sucrose overnight for cryoprotection.

The lumbar enlargement was cut into 50μm horizontal sections and sections containing the ventral horns were selected for immunohistochemistry. Detail of the antibodies can be found in supplementary table 1. Immunolabelling of AISs was made with an antibody against Ankyrin G (Rabbit anti-Ankyrin G, Santa Cruz, conc. 1:250), a scaffolding protein that spans the entire length of the AIS (Duflocq *et al*., 2011). To label motoneurones, an antibody against choline acetyltransferase (ChAT) was used (Goat anti-ChAT, Millipore conc. 1:100). Sections were blocked with 5% donkey serum in phosphate buffered saline with 0.3% triton X. Sections were incubated overnight in the primary antibodies diluted in the blocking solution then incubated sequentially with the secondary antibodies (donkey anti-rabbit Alexa Fluor 594 and donkey anti-goat Alexa Fluor 488, both Life technologies, conc. both 1:1000). Finally, the sections were mounted on glass slides and cover slipped using fluorescent mounting medium (Dako S3023).

Confocal stacks of the tissue were acquired using a Zeiss LSM 700 confocal microscope with a 20x objective. 6 sets of z-stacks in total were obtained per animal from lateral regions of ventral horn, these being motoneurones that innervate distal musculature. The selection criterion for AISs was that the entire AIS should be contained within the z-stack. ChAT staining was used to confirm that the AIS belonged to a motoneurone. For the cell body, the maximum and minimum diameters were measured to calculate the 2D surface area. For the AISs, the distal, the total length and the distance from the soma were measured in x, y, z planes using the ZEN Black 2.3 software (ZEISS).

Brightness and contrast were enhanced for the figures shown using Image J (NIH) which was performed uniformly to the whole image. All analysis, however, was performed on the raw data.

### Statistical analysis

All analysis was performed blinded up until the point of statistical tests which was performed using GraphPad Prism 7. The distribution of data was tested for normality using D’Agostino-Pearson omnibus normality tests and Brown-Forsythe tests (for significant differences in standard deviations) and parametric tests (ANOVA followed by Tukey’s post hoc tests) or non-parametric (Kruskall Wallis followed by Dunn’s multiple comparisons post hoc tests) were used accordingly. Data is shown as scatter dot plots with mean and standard deviation for parametric data or means and interquartile ranges for non-parametric data.

For data that co-varied with another value, linear regression was used for analysis and the differences between slopes were measured using Analysis of Covariance (ANCOVA). The level of significance for all the data was set to P <0.05. Full details of the statistical tests performed can be seen in tables 1 to 6.

**Table 1:** Details of statistical tests for data shown in figure 1

| Analysis | N (mice) | Medians (IQR) | Test | Test Statistic | P value | Post hoc test | Post hoc result |
| --- | --- | --- | --- | --- | --- | --- | --- |
| Rotarod (seconds) | YA: 14,<br>MA: 13<br>A: 16 | 192.2 (81)<br>128.7 (32.3)<br>79.0 (69.83) | Kruskal-Wallis | KW=26.44 | **$P<0.0001$** | Dunn's Multiple Comparisons | **YA vs MA: $P<0.05$**<br>**YA vs A: $P<0.0001$**<br>**MA vs A : $P<0.05$** |

**Table 2:**
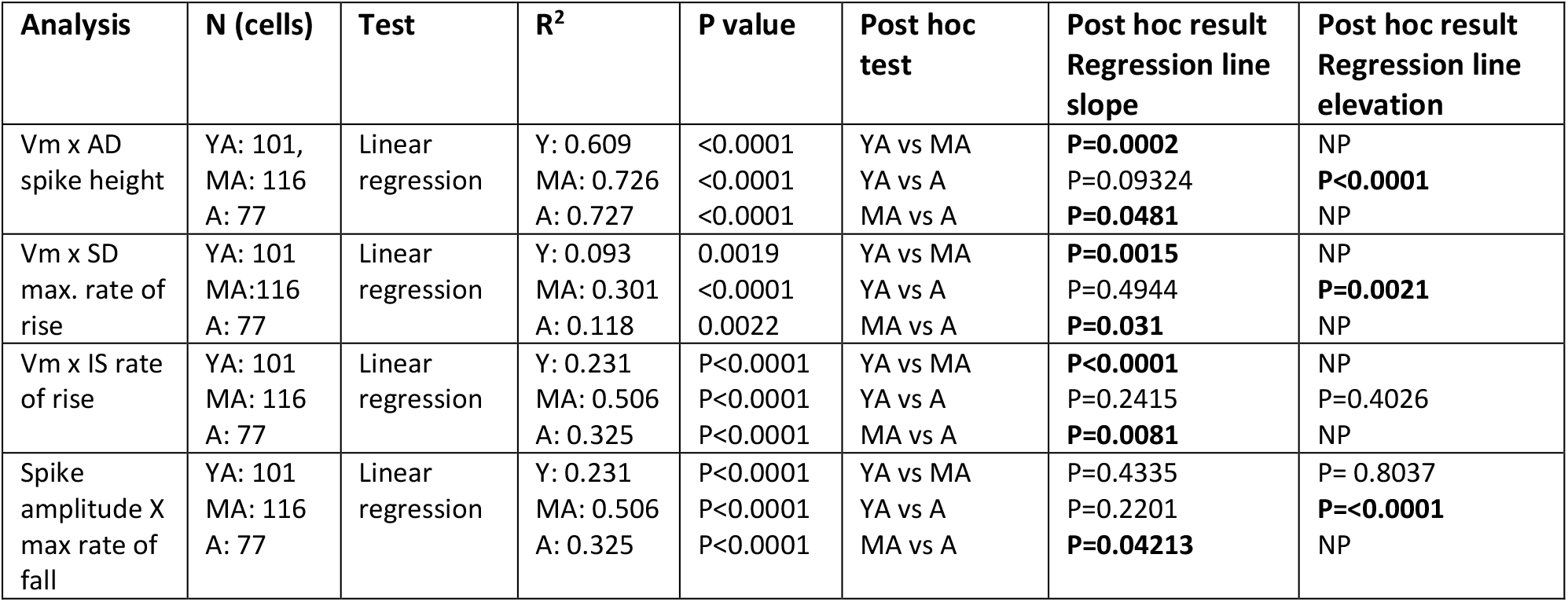
Details of statistical tests for linear regression shown in figure 2

| Analysis | N (cells) | Test | R<sup>2</sup> | P value | Post hoc test | Post hoc result Regression line slope | Post hoc result Regression line elevation |
| --- | --- | --- | --- | --- | --- | --- | --- |
| Vm x AD spike height | YA: 101, MA: 116 A: 77 | Linear regression | Y: 0.609<br>MA: 0.726<br>A: 0.727 | <0.0001<br><0.0001<br><0.0001 | YA vs MA<br>YA vs A<br>MA vs A | **P=0.0002**<br>P=0.09324<br>**P=0.0481** | NP<br>**P<0.0001**<br>NP |
| Vm x SD max. rate of rise | YA: 101 MA:116 A: 77 | Linear regression | Y: 0.093<br>MA: 0.301<br>A: 0.118 | 0.0019<br><0.0001<br>0.0022 | YA vs MA<br>YA vs A<br>MA vs A | **P=0.0015**<br>P=0.4944<br>**P=0.031** | NP<br>**P=0.0021**<br>NP |
| Vm x IS rate of rise | YA: 101 MA: 116 A: 77 | Linear regression | Y: 0.231<br>MA: 0.506<br>A: 0.325 | P<0.0001<br>P<0.0001<br>P<0.0001 | YA vs MA<br>YA vs A<br>MA vs A | **P<0.0001**<br>P=0.2415<br>**P=0.0081** | NP<br>P=0.4026<br>NP |
| Spike amplitude X max rate of fall | YA: 101 MA: 116 A: 77 | Linear regression | Y: 0.231<br>MA: 0.506<br>A: 0.325 | P<0.0001<br>P<0.0001<br>P<0.0001 | YA vs MA<br>YA vs A<br>MA vs A | P=0.4335<br>P=0.2201<br>**P=0.04213** | P= 0.8037<br>**P=<0.0001**<br>NP |

**Table 3:** Details of statistical tests for data shown in figure 2 (note linear regression data in table 2) *all data passed D’Agostino & Pearson omnibus normality tests and Brown-Forsythe test confirmed no significant differences in standard deviation. N.A = Not Applicable

| Analysis | N (cells unless otherwise stated) | Means (SD), Medians (IQR) or % | Test | Test Statistic | P value | Post hoc test | Post hoc result |
| --- | --- | --- | --- | --- | --- | --- | --- |
| SD spike width (2/3 amplitude ms) | YA:101 MA: 116 A: 77 | 0.2727 (0.0643)<br>0.2724 (0.0775)<br>0.2941 (0.0817) | Kruskal-Wallis | KW=12.71 | **P=0.0017** | Dunn's Multiple Comparisons | YA vs MA: NS<br>**YA vs A: P<0.01**<br>**MA vs A: P<0.01** |
| IS-SD delay (ms) | YA:101 MA: 116 A: 77 | 0.2772 (0.1008)<br>0.2520 (0.1008)<br>0.2520 (0.1392) | Kruskal-Wallis | KW=2.304 | P=0.3159 | N.A. | NA |

**Table 4:** Details of statistical tests for Axon Initial Segment (AIS) measurements (figure 3).

| Analysis | N | Medians (IQR) | Test | Test Statistic | P value | Post hoc test | Post hoc result |
| --- | --- | --- | --- | --- | --- | --- | --- |
| AIS length | YA: 336<br>MA: 335<br>A: 288 | 20.13 (5.45)<br>18.09 (7.19)<br>20.01 (6.13) | Kruskal-Wallis | KW=22.03 | < 0.0001 | Dunn's Multiple Comparison Test | **YA vs MA: P<0.0001**<br>YA vs A: NS<br>**MA vs A: P<0.01** |
| AIS distal width | YA:338<br>MA: 337<br>A: 290 | 1.29 (0.57)<br>1.05 (0.45)<br>1.29 (0.65) | Kruskal-Wallis | KW=46.76 | < 0.0001 | Dunn's Multiple Comparison Test | **YA vs MA: P<0.0001**<br>YA vs A: NS<br>**MA vs A: P<0.0001** |
| AIS distance from soma | YA:102<br>MA: 73<br>A: 77 | 7.53 (4.73)<br>7.19 (6.68)<br>5.93 (4.31) | Kruskal-Wallis | KW=3.815 | P=0.1485 | N.A. | N.A. |
| 2D soma area | YA: 103<br>MA:72<br>A:79 | 648.2 (200.8)<br>651.1 (300.9)<br>604.2 (160.5) | Kruskal-Wallis | KW=8.089 | P=0.0175 | Dunn's Multiple Comparison Test | YA vs MA: NS<br>**YA vs A: P<0.05**<br>**MA vs A: P<0.05** |

**Table 5:**
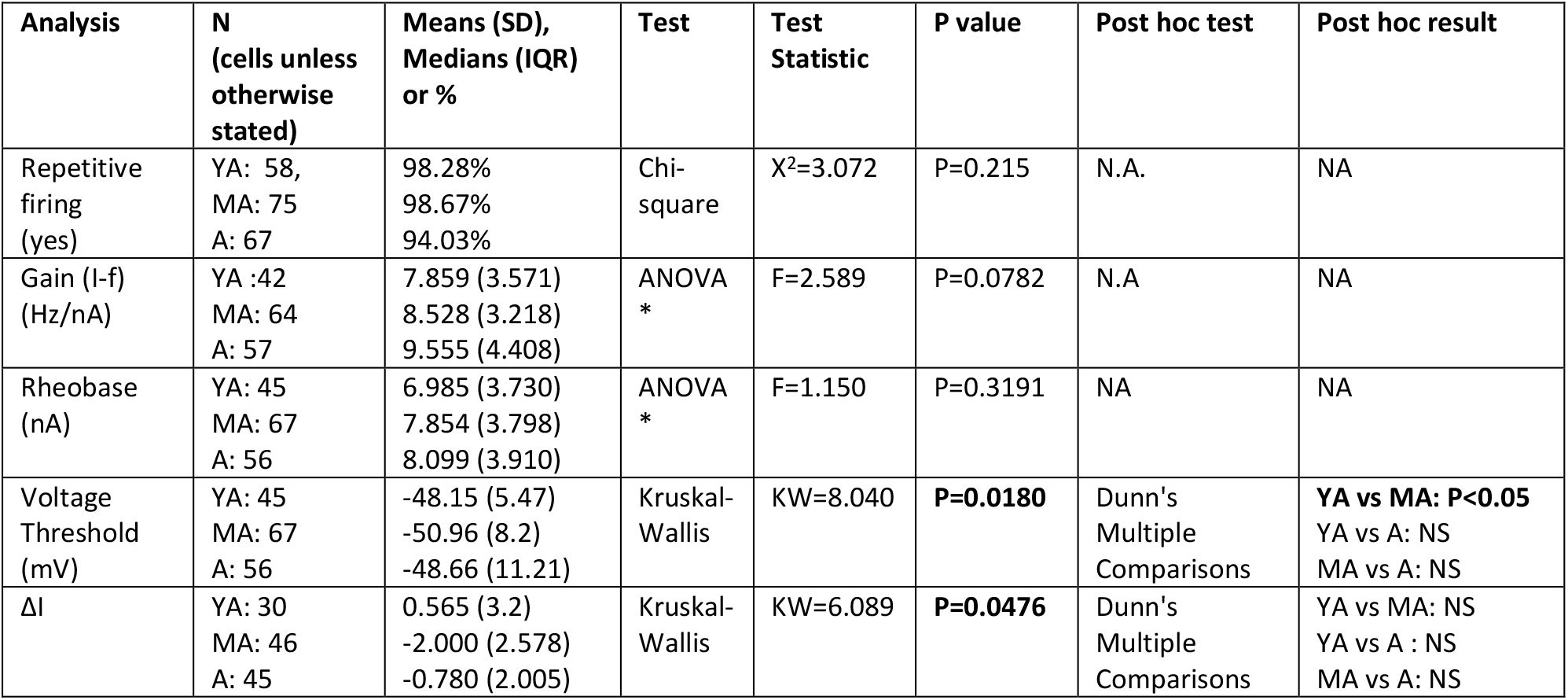
Details of statistical tests for data shown in figure 4 *all data passed D’Agostino & Pearson omnibus normality tests and Brown-Forsythe test confirmed no significant differences in standard deviation. N.A = Not Applicable

| Analysis | N<br>(cells unless<br>otherwise<br>stated) | Means (SD),<br>Medians (IQR)<br>or % | Test | Test<br>Statistic | P value | Post hoc test | Post hoc result |
| --- | --- | --- | --- | --- | --- | --- | --- |
| Repetitive<br>firing<br>(yes) | YA: 58,<br>MA: 75<br>A: 67 | 98.28%<br>98.67%<br>94.03% | Chi-square | $\chi^2=3.072$ | P=0.215 | N.A. | NA |
| Gain (I-f)<br>(Hz/nA) | YA :42<br>MA: 64<br>A: 57 | 7.859 (3.571)<br>8.528 (3.218)<br>9.555 (4.408) | ANOVA<br>* | F=2.589 | P=0.0782 | N.A | NA |
| Rheobase<br>(nA) | YA: 45<br>MA: 67<br>A: 56 | 6.985 (3.730)<br>7.854 (3.798)<br>8.099 (3.910) | ANOVA<br>* | F=1.150 | P=0.3191 | NA | NA |
| Voltage<br>Threshold<br>(mV) | YA: 45<br>MA: 67<br>A: 56 | -48.15 (5.47)<br>-50.96 (8.2)<br>-48.66 (11.21) | Kruskal-Wallis | KW=8.040 | **P=0.0180** | Dunn's<br>Multiple<br>Comparisons | **YA vs MA: P<0.05**<br>YA vs A: NS<br>MA vs A: NS |
| $\Delta I$ | YA: 30<br>MA: 46<br>A: 45 | 0.565 (3.2)<br>-2.000 (2.578)<br>-0.780 (2.005) | Kruskal-Wallis | KW=6.089 | **P=0.0476** | Dunn's<br>Multiple<br>Comparisons | YA vs MA: NS<br>YA vs A : NS<br>MA vs A: NS |

**Table 6:** Details of statistical tests for data shown in figure 5 and 6 for after-hyperpolarisation and input resistance N.A: not applicable

| Analysis | N (cells) | Medians (IQR) | Test | Test Statistic | P value | Post hoc test | Post hoc result |
| --- | --- | --- | --- | --- | --- | --- | --- |
| AHP Amplitude (mV) | YA:30<br>MA:64<br>A:57 | 1.33 (0.555)<br>1.69 (1.077)<br>1.61 (0.935) | Kruskal-Wallis | KW=7.545 | **P=0.023** | Dunn's Multiple Comparisons | **YA vs MA: P<0.05**<br>YA vs A: NS<br>MA vs A: NS |
| Vm for AHP (mV) | YA:30<br>MA:64<br>A:57 | -62.35 (18.54)<br>-63.71 (12.9)<br>-62.82 (18.15) | Kruskal-Wallis | KW=1.204 | P=0.5478 | N.A. | N.A. |
| AHP Duration ($^2/3$) | YA:30<br>MA:64<br>A:57 | 26.71 (12.94)<br>24.38 (5.74)<br>25.99 (8.22) | Kruskal-Wallis | KW=4.797 | P=0.0908 | N.A. | N.A. |
| Input Resistance (mΩ) | YA:57<br>MA:71<br>A:67 | 3.67 (2.15)<br>2.95 (1.74)<br>4.03 (1.93) | Kruskal-Wallis | KW=19.73 | **P<0.0001** | Dunn's Multiple Comparisons | **YA vs MA: P<0.05**<br>YA vs A: NS<br>**MA vs A: P<0.0001** |
| Absolute sag (mV) | YA:57<br>MA:71<br>A:67 | 0.788 (0.776)<br>0.820 (0.980)<br>0.651 (1.370) | Kruskal-Wallis | KW=2.231 | P=0.3278 | N.A. | N.A. |
| Relative sag (% total voltage drop) | YA:57:<br>MA:71<br>A:65 | 6.974 (7.963)<br>8.316 (11.075)<br>6.692 (9.877) | Kruskal-Wallis | KW= 7.610 | P=0.0223 | Dunn's Multiple Comparisons | YA vs MA: NS<br>YA vs A: NS<br>**MA vs A: P<0.05** |
| Proportion cells showing sag (%) | YA:57<br>MA:71<br>A:65 | 73.33%<br>83.58%<br>56.76% | Chi-square | X<sup>2</sup>= 12.5<br>df=2 | P=0.0019 | N.A. | N.A. |

## RESULTS

### Mice show a gradual decline in motor function

The motor function of 43 mice (YA:14, MA:13, A:16) was tested using the rotarod. The endurance time of the mice declined significantly with ageing (P<0.0001, Fig.1, Table 1). Post-hoc tests (Table 1) revealed significant differences between all groups confirming that the motor decline is gradual and already evident by middle-age in the mice. These mice were then used for either electrophysiological or anatomical experiments.

### Features of individual action potentials are altered by ageing

The electrophysiological data in these experiments come from 23 mice (YA: 8, MA: 9, A: 6). To start with, the features of single action potentials were analyzed. For this, averages of antidromic action potentials evoked from stimulation of the sciatic nerve were measured (YA:101, MA:116, A:77 action potentials) and representative examples are shown for each age group (Fig.2A). The amplitude of full antidromic action potentials correlated significantly with Vm for all groups (Fig.2B, Table 2). Both the angle and elevation of the slopes were then compared pairwise between groups. The angle of the slope was significantly different in the middle-aged mice compared to both the young adult and the aged mice (but not significantly different between the young and the aged mice). The elevation of the slope was significantly lower for the aged mice compared to the young adult, confirming that somatic action potentials are of lower amplitude in aged mice. The action potential width (measured at 2/3 amplitude) was, also, significantly wider in the aged mice than both young and middle-aged mice (Fig.2C, Table 3).

To determine if these alterations were due to a slower rate of rise or rate of fall of the action potential, the first derivative was obtained (see example in Fig.2D). From this the maximum rate of rise and fall could be obtained. From the example shown in Fig.2D it can be seen there is an initial fast rate of rise, shown as the first peak, which is then rapidly followed by a second peak. The first peak corresponds to the backfired action potential arriving at the axon initial segment (the IS component of the spike) and the second peak corresponds to the action potential at the soma-dendritic region (the SD component of the spike). To test the rate of rise of the spike at the soma we performed linear regression (and pairwise comparisons) for Vm by maximal rate of rise of the SD component of the action potential (Fig 2E). The regression slope for these were all significant (Table 2) but the elevation of this slope was significantly lower in the aged mice than the young adult mice confirming a slower rate of rise of somatic action potentials in the aged mice (Fig.2E, Table 2). As the rate of fall is heavily influenced by action potential amplitude, to investigate the rate of fall, regression slopes were made for maximum rate of fall by action potential amplitude (Fig.2F). The elevation of the regression slope was significantly lower in the aged mice than the young mice (Table 2) confirming a slower rate of fall of somatic action potentials in aged mice. A slower rate of rise and a slower rate of fall therefore both contribute to the increased width of somatic action potentials in aged mice.

These features so far, all measure aspects of the SD (somatic) portion of the antidromic action potential, therefore it can be concluded that the overall amplitude, rate of rise, rate of fall and width of somatic action potentials are altered in aged mice. To investigate the action potentials at the AIS the maximum rates of rise of the IS component of the antidromic action potentials were compared. The maximal rate of the rise of the IS component of the antidromic action potential correlated significantly with Vm for all groups (Fig.2G, Table 2), although, again, the angle of the regression slope for middle-aged mice was significantly different from both young and aged mice, having a much faster rate of rise at more hyperpolarized Vm (Fig.2G, Table 2). This increase was of a much larger magnitude than seen for the somatic portion of the antidromic spike. There was no significant difference between either the angle or the elevation of the regression slopes between young and aged mice (Fig.2G, Table 2). The faster rate of rise of the IS spike in middle-aged mice could be due to either an increased excitability at the AIS or the AIS being closer to the soma. However, the IS-SD delay measured as the time interval from the maximal rate of rise of the IS to the SD spike was not significantly different between groups (Fig.2H, Table 3).

### AISs of motoneurones in middle-aged mice show structural changes

To investigate if there are any structural changes to AISs that could explain the faster rate of rise of the IS spike in middle-aged mice, immunohistochemistry was used to label AISs using antibodies against the scaffolding protein Ankyrin G, a widely used marker of AISs (Fig.3A). Length, distal diameter and distance of the proximal end from soma were measured for AISs arising from ChAT immunoreactive motoneurones in the lateral column of the ventral spinal cord in 27 mice, 9 in each group. Detailed statistical analysis can be found in Table 4.

**Figure 3.**
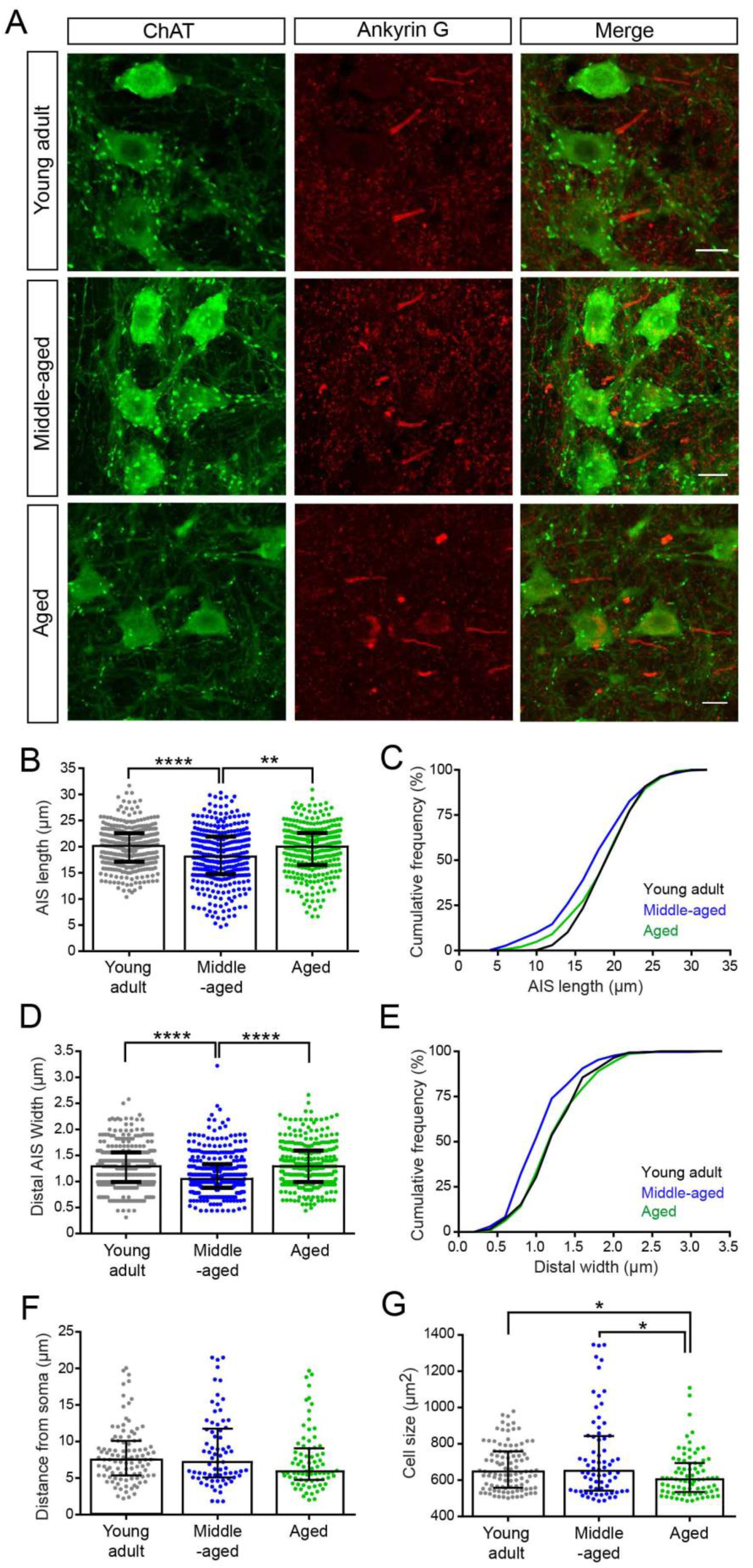
Axon initial segment parameters of motoneurones from mice at different ages. A. Representative examples of axon initial segments from YA, MA and aged mice. Motoneurones were identified using antibodies for Choline acetyl transferase (ChAT, in green) and axon initial segments with antibodies recognizing Ankyrin G (in red). Smaller clusters of ankyrin G labeling can also be seen as this is also found at nodes of Ranvier. Accumulation of Ankyrin G in the motoneurone cell bodies was commonly observed in the aged mice. Scale bar 20μm. B. Scattered dot plot of axon initial segment length (μm) from young adult (n=336 cells), middle aged (n=335 cells) and aged (n=288 cells) mice. The axon initial segments in the MA mice are significantly shorter compared to the YA and aged mice. C. Cumulative frequency distribution of the axon initial segment length (μm) from YA, MA and aged mice showing that a large percentage of MA mice have shorter axon initial segments. D. Scattered dot plot of distal width (μm) of motoneurone axon initial segments from YA (n=338 cells), MA (n=337 cells) and aged (n=290 cells) mice. The axon initial segments of the MA mice are significantly thinner compared to the YA and aged mice. E. Cumulative frequency distribution of the axon initial segment distal width (μm) from YA, MA and aged mice showing that a large percentage of MA mice have thinner initial segments. F. Scattered dot plot of axon initial segment distance from the motoneurone soma (μm) from YA (n=102 cells), MA (n=73 cells) and aged (n=77 cells) mice. No significant differences across the different age groups were found. G. Scattered dot plot of motoneurone cell size (μm^2^) from YA (n=103 cell), MA (n=72 cells) and aged (n=79 cells) mice. The motoneurone cell size in the aged mice is significantly smaller compared to the YA and MA mice. Data presented as individual data points with median (+interquartile ranges), *P<0.05, **P<0.01, ***P<0.001 and ****P<0.0001 by Kruskal-Wallis test.

Statistical analysis of these measurements revealed that the median AIS length in the middle-aged mice was approximately 10% shorter than the median AIS length in both young adult and aged mice. There was no difference in AIS length between the young adult and aged mice (Fig.3B and C, Table 4). The median AIS diameter (measured at the distal end) was also 18.6% thinner in the middle-aged mice than both the young adult and aged mice (Fig.3D and E, Table 4).

Although the median distance of the AIS from the soma in aged mice was less than in young adult and middle age mice respectively, this was not significant (Fig.3F, Table 4). Distance from soma therefore cannot explain the faster rate of rise seen in the IS component of the antidromic spike in the middle-aged mice. In addition, there was also no difference between young adult and middleaged mice with respect to distance of the AIS from the soma, also, ruling out this explanation. The decrease in the AIS diameter observed in the middle-aged mice could potentially account for the increased rate of rise of the AIS spike. A lower diameter axon initial segment would presumably be more excitable if this represented a higher density of Na^+^ channels in a smaller surface area.

As AIS parameters are influenced by cell size, we therefore also measured soma size. The average soma area of motoneurones in middle-aged mice was similar to the young mice. In the aged group motoneurones, however, were significantly smaller than both the young and middle-aged group (Fig.3G, Table 4).

### Features of repetitive firing are unaltered during ageing

AIS length is an important regulator for modulating the repetitive firing in neurones (Evans *et al*., 2015). Therefore, to test for the effects on repetitive firing behaviour, triangular current injections were used to evoke repetitive firing in the motoneurones, examples of which are illustrated in Fig. 4A. 57/58 young adult, 74/75 middle-aged and 63/67 aged motoneurones responded with repetitive firing. These proportions were not significantly different between groups (Fig.4B, Table 5) confirming no difference in the ability to produce repetitive firing with ageing (in fact 3 of the 4 cells not firing in aged mice came from the same mouse using the same electrode).

**Figure 4.**
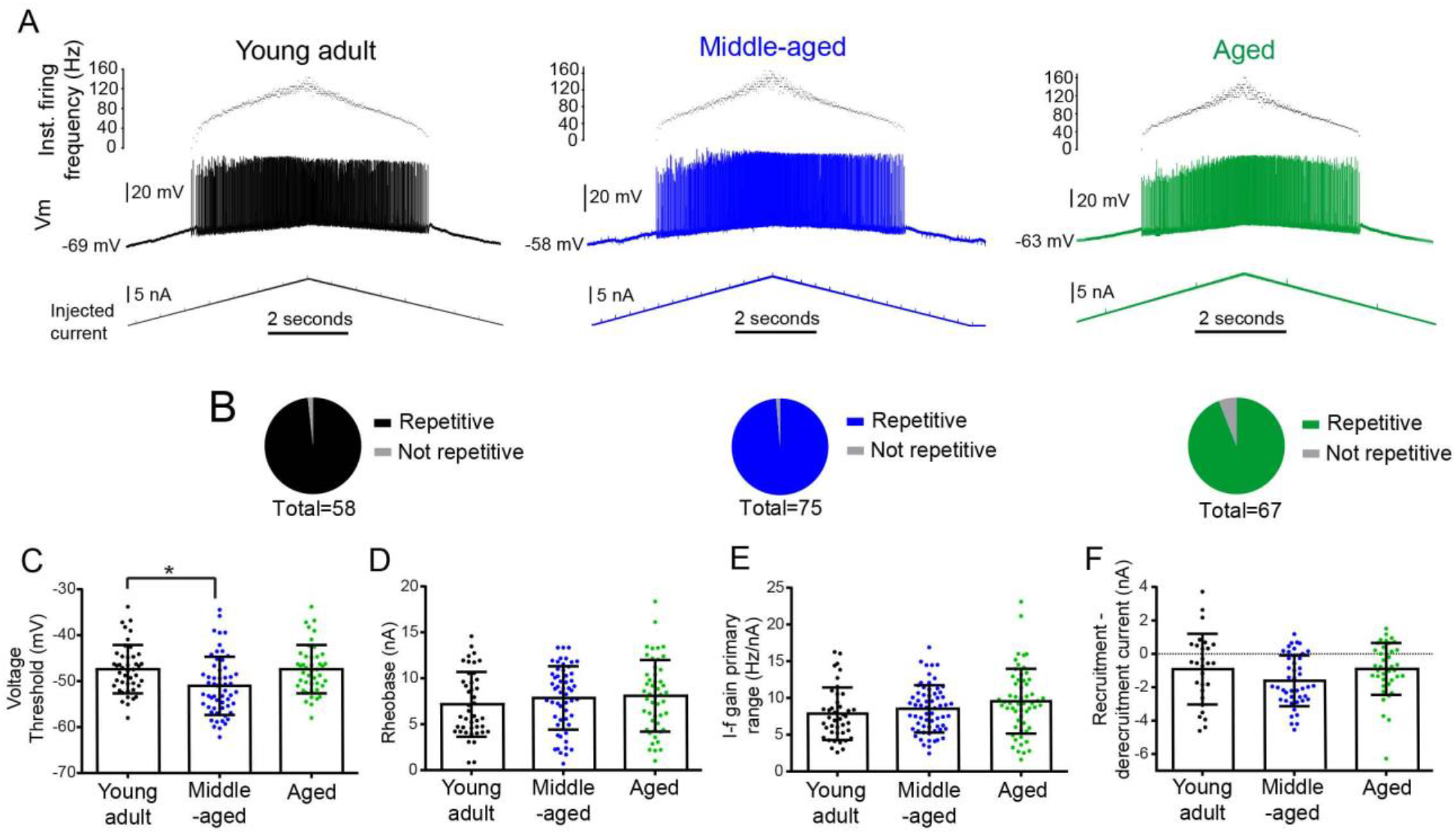
Features of repetitive firing in motoneurones in response to intracellular current injection in Young Adult (YA), Middle-Aged (MA) and aged mice. A. Examples of repetitive firing recorded intracellularly (middle traces) in motoneurones from YA (black), MA (blue) and aged (green) mice in response to intracellular injection of ramps of ascending and descending intensity of current (bottom trace). The top trace shows the instantaneous firing frequency. B. The proportions of motoneurones from YA (black), MA (blue) and aged (green) mice, responding to the current injection with repetitive firing which was not significantly different (n = 58 cells in YA, 75 cells in MA and 67 cells in aged mice). C. Scatter dot plot showing data points for individual cells for the voltage thresholds at which repetitive firing was evoked in motoneurones, which was at significantly more hyperpolarized values in the MA mice ( n = 45 cells in YA, 67 cells in MA and 56 cells in aged mice). Data is shown with median (+interquartile ranges). D. Scatter dot plot showing data points for individual cells for the rheobase currents at which repetitive firing was evoked in motoneurones, which was not significantly different between groups. Data is shown with means (and standard deviations). E. Scatter dot plot showing data points for individual cells for I-f gain in the primary range for cells from YA, MA and aged mice which was not significantly different between groups. Data is shown with mean (+SD), n = 42 cells in YA, 64 cells in MA and 57 cells in aged mice. F. Scatter dot plot showing individual data points with means and SD for recruitment – de-recruitment currents for motoneurones from YA, MA and aged mice. Data is shown with median (+interquartile ranges) and shows a trend for decreased PIC activation in MA mice (n = 30 cells in YA, 46 cells in MA and 45 cells in aged mice).

As the initial rate of rise of the IS component on the action potential was faster in the middle-aged mice, one could predict that this would make it easier to reach threshold for repetitive firing. Consistent with this, the voltage threshold for repetitive firing measured in the soma was significantly lower in the middle-aged mice than both young adult and aged mice (Fig.4C, Table 5). Surprisingly, however, rheobase currents for repetitive firing were not significantly different between any of the groups (Fig.4D, Table 5). Similarly, the gain of the motoneurone which was obtained using current-frequency (I-f) slopes and measured during the primary range of firing was, also, not significantly different between any of the groups (Fig.4E, Table 5). This is not what would be expected with a more excitable AIS.

Persistent inward currents and/or spike frequency adaption can also influence the I-f slope, a crude measure of which is to subtract the de-recruitment current from the recruitment current (ΔI). A higher value is normally used to indicate higher levels of persistent inward current. In these experiments we saw that this value decreases slightly in the middle-aged mice (Figure 4F, Table 5), suggesting decreased persistent inward current or enhanced spike frequency adaptation at this age. The I-f gain of motoneurone is, also, heavily influenced by the post-spike after-hyperpolarization, therefore we investigated this next.

### The after-hyperpolarization amplitude but not duration is altered during ageing

Brief current pulses were used to evoke orthodromic action potentials in the motoneurones (YA:30, MA:64, A:57 cells). An example of how AHPs were measured at 2/3 recovery is shown in Fig.5A and B. AHPs from middle-aged mice had significantly larger amplitudes than young mice (Fig.5C, Table 6). This was not due to differences in resting Vm as this was not significantly different between groups (Table 6). However, ageing did not appear to have any effect on AHP duration (Fig.5D, Table 6).

**Figure 5.**
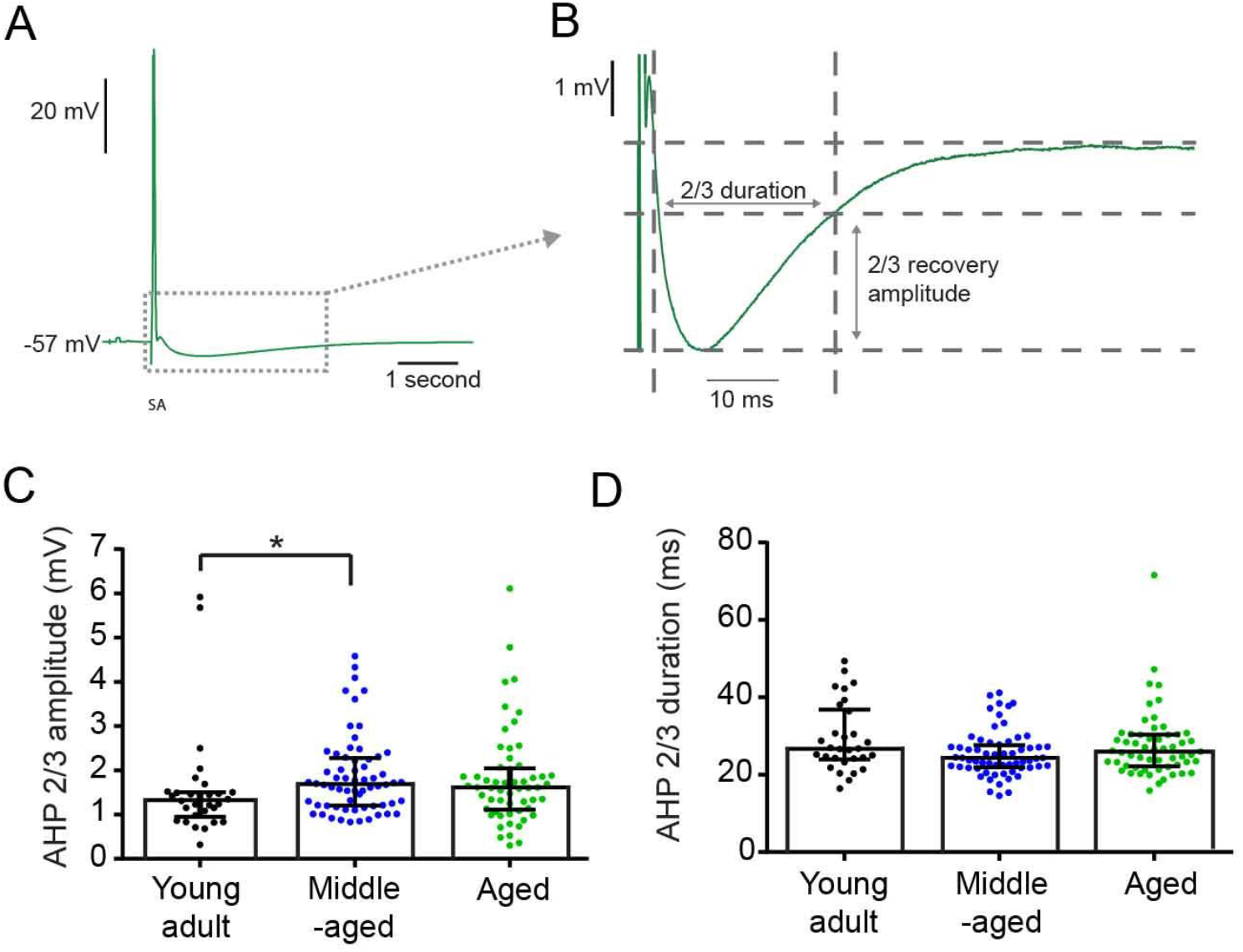
Features of After-hyperpolarizations (AHPs) of action potentials in motoneurones from mice at different ages. A. Example of an action potential evoked by a brief current pulse in an aged motoneurone. B. Details of the measurement of amplitude and duration both measured at 2/3 relaxation time. C. Scatter dot plot showing individual data points for AHP amplitude for action potentials from YA (black), MA (blue) and aged (green) mice. This was significantly different between YA and MA mice. Data is shown with median (+interquartile ranges). D. Scatter dot plot showing individual data points for AHP duration (measured at 2/3 amplitude) for action potentials from YA (black), MA (blue) and aged (green) mice. Data is shown with median (+interquartile ranges) which were not significantly different between groups.

### Input resistance is reduced in middle-aged mice

Differences in somatic input resistance could also cancel out the effects of a more excitable axon initial segment. This was measured in 195 motoneurones (YA:57, MA:71, A:67) using the voltage response to a −3nA hyperpolarizing current, examples of which are shown in Fig.6A. Significant differences were found with respect to age and post-hoc tests showing this to be mainly due to a reduction in input resistance occurring at middle age, which returned to normal values at 600 days (Fig.6B, Table 6).

**Figure 6.**
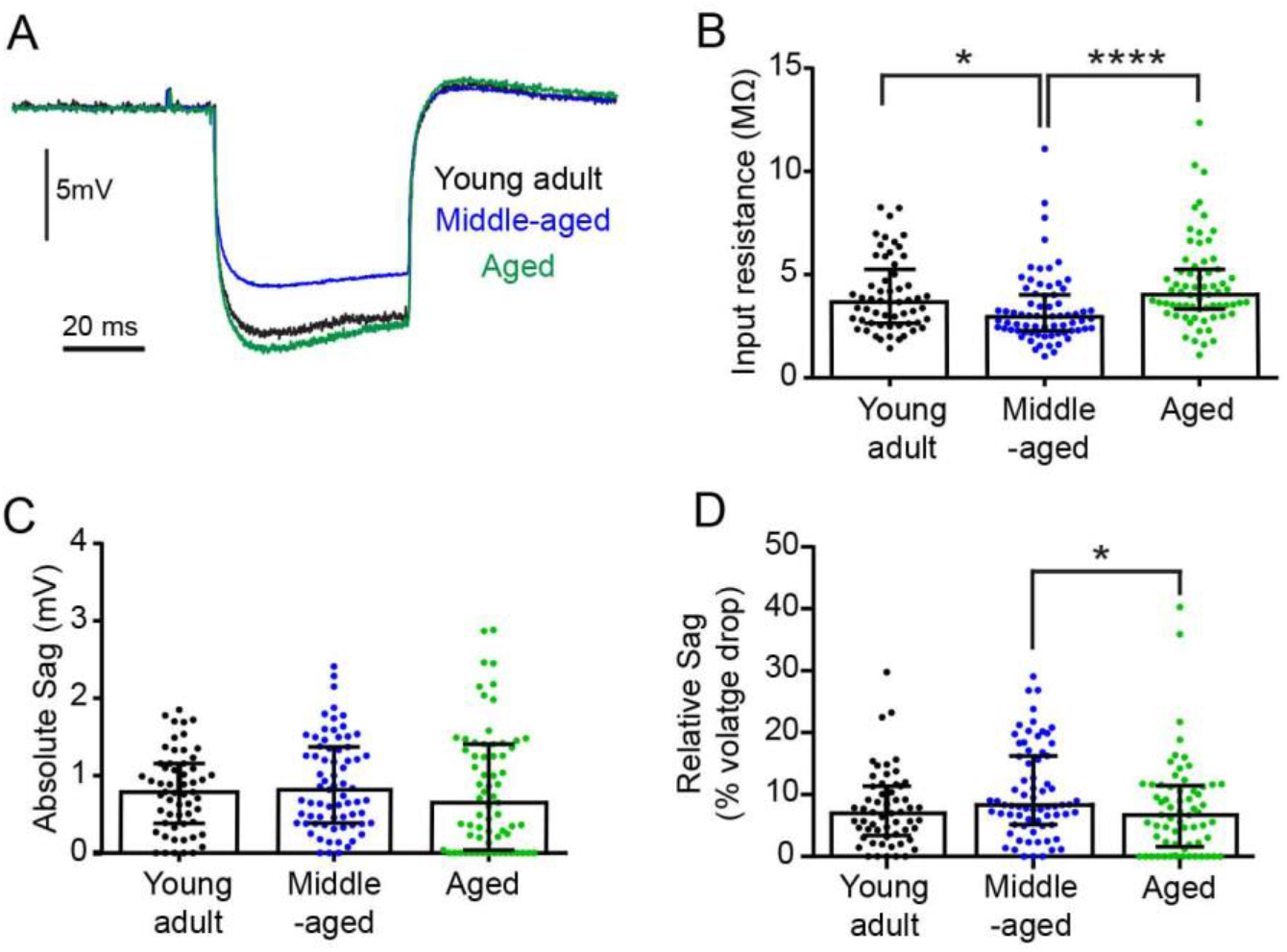
Input resistance changes with ageing. A. Examples of the averages of the voltage drop in response to intracellular injections of 50ms, −3nA hyperpolarising square current pulses from motoneurones in young adult (YA), middle aged (MA) and aged (A) mice. B. Scatter dot plot showing the input resistances for individual cells calculated from the peak voltage drop, which was significantly lower in MA mice than both YA and aged mice. C. Scatter dot plot showing the sag recorded at the end of the 50 ms long −3nA current pulse which was not significantly different between groups. D. Scatter dot plot for individual cells showing the same sag data as C but expressed as a percentage of the total voltage drop, which was significantly different between MA and aged mice. Data presented as individual data points with median (+interquartile ranges). For all analysis n= 57 cells YA, 71 cells MA and 65 cells in aged mice

Many of the voltage responses showed a delayed sag (indicated on Fig.6A) which has been previously demonstrated in mouse motoneurones (Manuel *et al*., 2009;Meehan *et al*., 2010b). The sag has been attributed to the activation of Hyperpolarization-activated Cyclic Nucleotide–gated (HCN) channels (Ito & Oshima, 1965). The absolute sag measured at the end of the 50 ms hyperpolarizing current pulse was not significantly different between age groups (Fig. 6B, Table 6). To control for differences in the input resistance the sag was also expressed as a percentage of the peak (maximum) voltage drop. This became significantly larger in middle-aged mice and then decreased again in aged mice (although the difference only reached significance with aged mice, Fig.6C, Table 6). Together, these results suggest that hyperpolarization-activated current (Ih) currents are slightly increased and compensating for the decreased input resistance at middle age. Although the proportional magnitude of the sag appears to normalize towards the original levels in the aged mice it should be noticed that a larger proportion of cells in the aged mice showed no sag (Table 6).

## DISCUSSION

Our results clearly demonstrate that the intrinsic electrical properties of spinal motoneurones change not just with advanced ageing but throughout the lifetime of mice. What is perhaps most surprising is that the changes are not linear and, in some cases, even reverse after middle age a scan be seen in Figure 7 where the changes in the different parameters can be seen relative to one another. Even more surprisingly, we observed that the basic gain of the motoneurone (in terms of the input-firing frequency output relationship) was relatively unchanged even in the aged mice. This suggests that there are homeostatic mechanisms in play, which are very effective in maintaining normal output. Furthermore, this suggests that it is unlikely that impairments in the ability of the soma/AIS to initiate repetitive firing in response to input contribute to the age-related decline in motor function.

**Figure 7:**
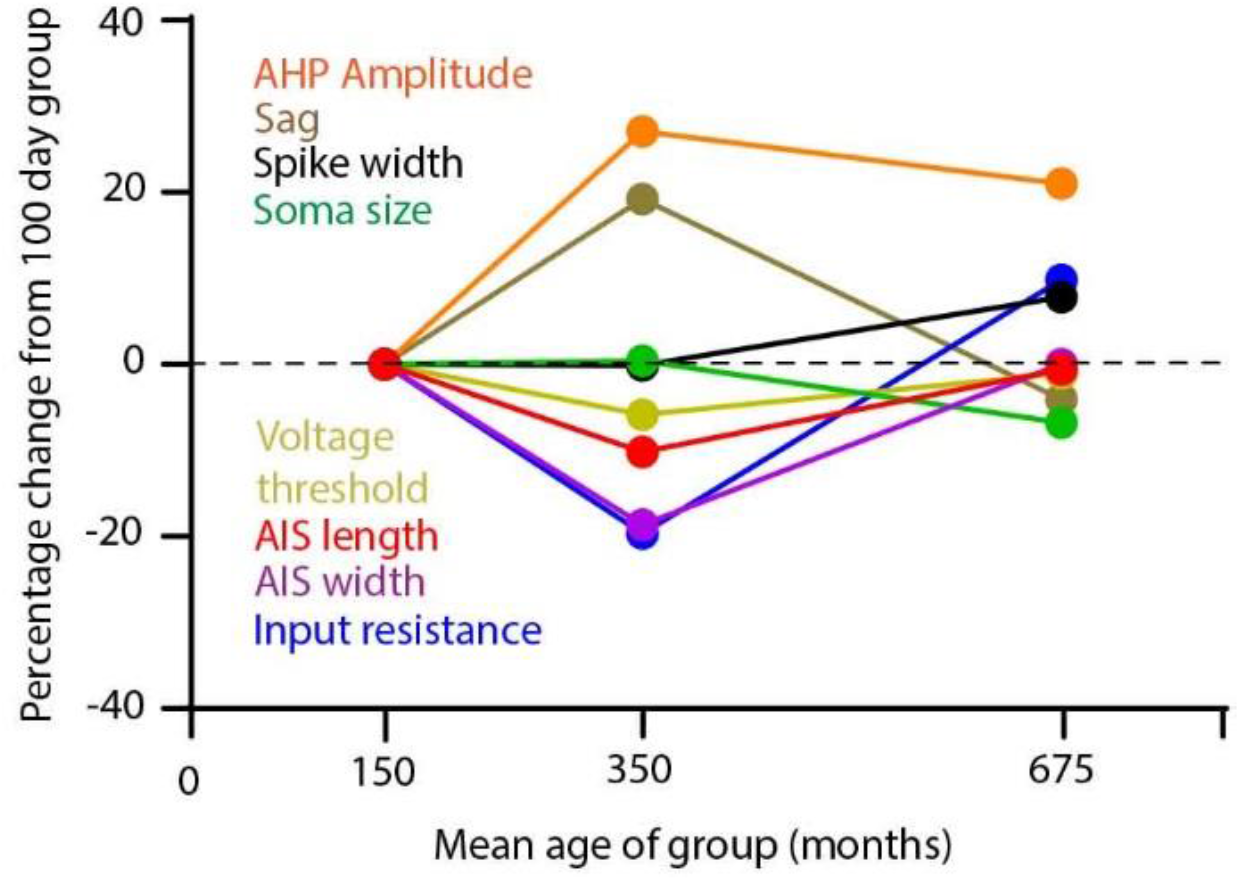
Changes in intrinsic properties of spinal motoneurones over the lifetime of the mouse. Graph showing how the various parameters measured change with age relative to one another normalized to the measurements at 100 days old

### Implications for ageing

Recent studies have found surprisingly small or no reductions in the amplitude of compound muscle action potentials from sciatic nerve stimulation in aged mice at a similar age to that used here (Giorgetti *et al*., 2019;Sheth *et al*., 2018). However, both studies using a Motor Unit Number Estimation (MUNE) method elegantly demonstrated a loss of functional motor units. Our results suggest that it is highly unlikely that inability of the motoneurones to produce functional output can account for the reduction in the number of functional motor units observed using MUNE.

One logical explanation for the apparent loss of motor units seen with MUNE could be an age-related motoneurone death, although this appears to be a highly contentious issue. Early postmortem studies in humans revealed that the number of lumbar motoneurones remains relatively stable up until 60 years of age, which is followed by a gradual loss of up to 25% by 90 years of age (Cruz-Sanchez *et al*., 1998;Kawamura *et al*., 1977;Tomlinson & Irving, 1977). Furthermore, some studies show an age-related decrease in the number of lumbar motoneurones in rats (Jacob, 1998;Hashizume *et al*., 1988) and in C57BL/6J mice (Piekarz *et al*., 2020), while others have found no significant loss of motoneurones in aged cats (Liu *et al*., 1996), rats (Chopek & Gardiner, 2010) or C57BL/6J mice at 28-29 months (Chai *et al*., 2011;Maxwell *et al*., 2018). Interestingly, Chai et al. reported a drastic denervation of hind limb muscles of the mice at the same age (Chai *et al*., 2011), which would account for the decrease in functional motor units.

#### Declines in persistent inward currents would make the motoneurones less responsive to synaptic input

In our experiments, we mimic synaptic input by direct current injection to the soma. This would activate the axon initial segment directly to evoke repetitive firing without the influence of the dendritic ion channels. Thus, our measurements of the rheobase, voltage threshold and primary range of firing only reflect excitability of the soma-AIS regions. At higher current levels, the depolarization reaches the dendritic regions where the L-type calcium channels mediating persistent inward currents are located. Using the dI method we were able to detect a trend for some alterations indicating a decreased activation of persistent inward currents as mice reached middle age. This fits partially with the observations in humans using an indirect version of this technique (based on voluntary triangular force contractions) which indicated a decrease in persistent inward currents in aged (~70 year old) humans (Hassan *et al*., 2021; Orssatto *et al*., 2021) except that we see this in middle age with a restoration in the much older mice. Although we must express caution here in comparison as the experiments in humans are performed in awake conscious subjects compared with anaesthetized mice where persistent inward currents can be suppressed either by a direct effect on the ion channels or due to a reduction in monoaminergic modulation of these channels. The secondary range of firing, normally associated with activation of dendritic persistent inward currents was notably less frequent and less obvious in our current experiments than we normally observe when using our preferred Hypnorm and Midazolam aneasthesia. However, Hypnorm was unavailable at the time of the recordings due to a pause in production. Thus, a more accurate analyses of persistent inward currents would be best achieved in an unanaesthetised decerebrate preparation (Meehan *et al*., 2017).

#### Increased input resistance is a consistent feature of aged motoneurones across species

Another implication of our observations of non-linear changes in excitability throughout the lifetime of a motoneurone is that it makes comparison between studies difficult, particularly those across species. This leaves the pertinent question: what ages should be considered as a “control” group against which to compare results obtained in aged animals? Clearly our conclusions regarding what constitutes increases or decreases in certain parameters would depend on whether we compare the aged results with young adult or middle-aged animals. This emphasizes the importance of examining changes in old age in the context of the ongoing changes occurring across the lifetime.

One consistent observation across cat and rat has been the increase in input resistance in advanced age (Kalmar *et al*., 2009;Morales *et al*., 1987). While here we observed decreased input resistance at middle age in mice, this did increase again in the aged animals. Analysis of changes in membrane time constants and total cell capacitance from the recordings in the cat suggested that the increased input resistance with ageing is due to both increases in membrane resistance and decreases in soma surface area (Engelhardt *et al*., 1989). This latter feature has been confirmed by anatomical finding of decreases in soma size in cat (Liu *et al*., 1996), rat (Hashizume *et al*., 1988) and in the aged mouse, as we have demonstrated here. Interestingly, a recent study in monkey and mouse did not find reductions in the size of the motoneurone somas (Maxwell *et al*., 2018). In that study, alpha motoneurones were identified using Neu-N and C-bouton labelling which itself is reduced with ageing (Maxwell *et al*., 2018). We specifically used ChAT labelling to identify motoneurones and restricted our analysis to those motoneurones in the lateral motoneurone columns at the segment corresponding to the known location of hind limb motoneurones. We therefore are confident that these represent the same population from which we were recording from in the electrophysiology experiments.

AHP duration has been estimated in humans using the firing frequency of motor units recorded with electromyography which has indicated that AHP duration increases with age (Piotrkiewicz *et al*., 2007). Recordings in rats, also, show an increase in both AHP amplitude and duration with age (Kalmar *et al*., 2009). Our results, however, showed no changes in AHP duration with age which is consistent with the results in cat (Morales *et al*., 1987). These differences are unlikely to be due to anesthesia as we used the same ketamine-xylazine anesthetics in our mice as was used in the rats (Kalmar *et al*., 2009), suggesting that changes in AHP duration are not straightforward and may require further, more comprehensive investigation. We, also, failed to observe significant difference in rheobase in the aged mice as has been seen in cats and rats raising the possibility that not all effects of ageing are uniform across species. This is important to acknowledge, since modern research on neurodegenerative diseases affecting the motoneurone heavily relies on transgenic mouse models and this must be taken into account when comparing changes seen in such models with healthy ageing.

### Homeostasis masks the motoneurone “mid-life crisis”

During middle age in mice we observed changes in the intrinsic properties of spinal motoneurones, which are highly unlikely to be due to selective survival of a certain motoneurone population or due to loss of functional motor units as previously shown with MUNE methods for middle-aged mice (Sheth *et al*., 2018). Furthermore, many of these factors return to normal with advancing age, as we have shown in the aged mice. Therefore, this indicates that there are homeostatic mechanisms which act throughout the lifetime, and the changes that we observed represent true plasticity of the motoneurones.

The decrease in input resistance we observed should make it harder to reach the voltage threshold for action potentials. But, if constriction of the AIS increases the relative density of Na^+^ channels (an assumption consistent with an increased rate of rise of the IS portion of the antidromic spikes) then this could increase the excitability of the AIS and compensate for the reduction in somatic input resistance. This could explain why we do not see significant changes in rheobase, despite the altered input resistance. This, along with the reductions in persistent inward currents would also maintain a normal I-f gain. If this is the case, then one could predict that as the voltage threshold is measured with the microelectrode in the soma and not directly at the AIS, then the imbalance in excitability between the AIS and the soma would appear as a lower voltage threshold recorded at the soma. This is precisely what we observed. Therefore, we conclude that changes at the AIS most likely represent a homeostatic response to counteract soma-dendritic changes that occur around middle age that reduce input resistance.

### There is no one way for an axon initial segment to age

Our results add to the growing knowledge on AIS changes in ageing. Initial reports investigating AIS plasticity in aged marmoset monkeys found AIS shortening across a wide range of brain regions including frontal, prefrontal and occipital regions (Atapour & Rosa, 2017). Similar observations were made for the AISs of neurones located in the primary visual cortex in aged rats (Ding *et al*., 2018). This suggested that AIS shortening might be a general feature of ageing. However, in the hippocampus, significant shortening of AISs was not found across different regions in middle-aged or aged rats (Kneynsberg & Kanaan, 2017). While we did observe AIS shortening during middle age on the mouse motoneurones this seemed to be a transient change, which returned to previous levels in the aged mice. Thus, our results support the view that AIS shortening is not a global change across the nervous system in old age.

This is perhaps intuitive if the changes are truly homeostatic rather than degenerative in nature as some neuronal types in the CNS will experience drastically different age-related perturbation in inputs. Furthermore, modelling studies report that optimal AIS length is dependent on the size of the soma and the dendritic region, therefore the same AIS changes may have different consequences for neuronal excitability in different cells (Gulledge & Bravo, 2016). For example, in small cells with short AISs, increases in AIS length causes an increase in rheobase in contrast to larger cells (such as motoneurones) where a decrease in rheobase is seen. It is also possible that modifications of the AIS geometry may reflect changes in the entire soma-dendritic region occurring with ageing. Although, to prove this requires technically challenging intracellular recording and subsequent painstaking cell reconstruction. The only such study to date revealed that the dendritic trees of motoneurones innervating intrinsic foot muscles had much more extensive branching in aged cats (Ramirez & Ulfhake, 1992).

In addition to changes in length we also observed changes in AIS diameter with ageing. The published data on AISs on hippocampal neurones in aged rats (Kneynsberg & Kanaan, 2017) is the only other study to investigate possible changes in diameter (presumably due to the difficulty in measuring AISs which are typically much thinner compared to the AISs of spinal motoneurones). According to this rat study, significant decreases in AIS diameter were observed only in the CA2 region, the same region where the decrease in AIS length was observed in the aged rats, with a similar trend seen in the middle-aged rats (Kneynsberg & Kanaan, 2017). This is in agreement with our observations in middle-aged mice, although both the length and diameter returned to normal values in the aged mice. Studies of AIS plasticity, including modeling, tend to pay little attention to changes in AIS diameter, focusing on length and location instead, however, changes in diameter may be a very efficient way of altering the excitability of the AIS. According to mathematical models of the node of Ranvier, constriction of the node can significantly increase nodal excitability and increase conduction velocity in axons (Johnson *et al*., 2015; Halter & Clark, Jr., 1993). Likewise, constriction of the AIS, if it serves to increase the density of Na^+^ channels, might also result in increased excitability of the AIS. Of course, we do not know if this is the case here, however, our observations of a faster rate of rise of the IS component of the antidromic action potentials would surely be consistent with this hypothesis.

### Homeostasis in ageing and disease: Comparisons with excitability changes in neurodegenerative motoneurone diseases

Ageing is one of the key elements in a number of pathological cascades leading to development of large number of CNS disorders, therefore, it is crucial to have a comprehensive understanding on the effects that advancing age has on the neuronal homeostasis. Our observations of homeostatic changes occurring in middle age are particularly intriguing in the context of neurodegenerative diseases affecting motoneurones that tend to have an adult onset (from middle age), such as Amyotrophic Lateral Sclerosis (ALS). Excitability changes have been observed in our laboratory and others using mouse models of this diseases and it is relevant to compare these with the changes we have observed to determine if they represent exacerbated forms of ageing or a failure of the homeostatic mechanisms, an interesting concept that has recently been suggested (Kuo *et al*., 2020)

Immediately prior to symptom onset in the G127X SOD1 mouse model of ALS we have observed shorter AISs on spinal motoneurones (Bonnevie *et al*., 2020) which then elongates but becomes thinner as symptoms onset (Jørgensen HS *et al*., 2020). This is, also, accompanied by increased rate of rise of the IS component of the antidromic action potential, hence AIS constriction in symptomatic mice (~225 days old) appears to mirror the changes observed in the middle-aged mice but occurring earlier. Although it should be noted that the eventual later increase in AIS length seen in the symptomatic G127X SOD1 mice actually exceeds the maximum length observed at any time points in these mice. However, this may be a homeostatic response to control for the loss of descending motor drive which is much more pronounced in this disease than in normal ageing.

Another prominent feature of the ALS SOD1 mouse models is an increase in persistent inward currents, both Ca^2+^ and Na^+^, a feature already present in early embryonic/neonatal stage (Kuo *et al*., 2005;Quinlan *et al*., 2011) and persisting into adulthood (Bonnevie *et al*., 2020;Delestree *et al*., 2014;Huh *et al*., 2020;Jensen *et al*., 2020a; Jørgensen HS *et al*., 2020;Meehan *et al*., 2010a). In the current study, as the C57Bl/6J mice reach middle age, we actually see the opposite, with indications of decreases in persistent inward currents (although it should be acknowledged that the ΔI method is an indirect measure). It has been suggested that this increased persistent inward currents compensate for increases in input conductance and change in parallel with this as the disease progresses (Huh *et al*., 2020). It therefore appears that increased persistent inward current in ALS is a distinct pathophysiological feature of this disease rather than an exacerbation of the normal ageing process in motoneurones.

## Conclusions

A cascade of homeostatic changes occur during the lifetime of a spinal motoneurone to ensure a remarkably constant physiologically functional output response to input. The most extreme excitability changes seen in motoneurones in neurodegenerative diseases do not appear to be simply exacerbated version of these changes and are most likely in response to other pathological alterations occurring during the disease process.

## Acknowledgments

We acknowledge the Core Facility for Integrated Microscopy, Faculty of Health and Medical Sciences, University of Copenhagen.

## Notes

**Funding Sources**, This work was supported by project grants from the Lundbeck foundation.

### Competing Interest Statement

The authors have declared no competing interest.

